# SARS-CoV-2 Omicron BA.2.87.1 Exhibits Higher Susceptibility to Serum Neutralization Than EG.5.1 and JN.1

**DOI:** 10.1101/2024.03.10.584306

**Authors:** Qian Wang, Yicheng Guo, Logan T. Schwanz, Ian A. Mellis, Yiwei Sun, Yiming Qu, Guillaume Urtecho, Riccardo Valdez, Emily Stoneman, Aubree Gordon, Harris H. Wang, Lihong Liu, David D. Ho

## Abstract

As SARS-CoV-2 continues to spread and mutate, tracking the viral evolutionary trajectory and understanding the functional consequences of its mutations remain crucial. Here, we characterized the antibody evasion, ACE2 receptor engagement, and viral infectivity of the highly mutated SARS-CoV-2 Omicron subvariant BA.2.87.1. Compared with other Omicron subvariants, including EG.5.1 and the current predominant JN.1, BA.2.87.1 exhibits less immune evasion, reduced viral receptor engagement, and comparable infectivity in Calu-3 lung cells. Intriguingly, two large deletions (Δ15-26 and Δ136-146) in the N-terminal domain (NTD) of the spike protein facilitate subtly increased antibody evasion but significantly diminish viral infectivity. Collectively, our data support the announcement by the USA CDC that the public health risk posed by BA.2.87.1 appears to be low.

## Main text

SARS-CoV-2 Omicron subvariant BA.2.87.1, which emerged in South Africa in September 2023^1^, has garnered global interest due to its unique constellation of many mutations in the spike protein. The highly mutated BA.2.87.1 is derived from BA.2, and among other changes, it features two large deletions (Δ15-26 and Δ136-146) in the N-terminal domain (NTD) of the spike protein not observed in any other variants, including Omicron BA.2, EG.5.1, or the currently dominant JN.1 **(Figure 1A)**. These deletions, along with an additional substitution, W152L, are located near the NTD antigenic supersite that is targeted by many neutralizing antibodies (**Figure S1)**. Consequently, investigating whether the novel variant BA.2.87.1 has enhanced immune evasion and infectivity is important, as it may pose a potential threat to public health. Furthermore, characterization of BA.2.87.1 and its uniquely large NTD deletions may contribute to understanding the limits of functional SARS-CoV-2 spike protein changes.

**Figure 1.**
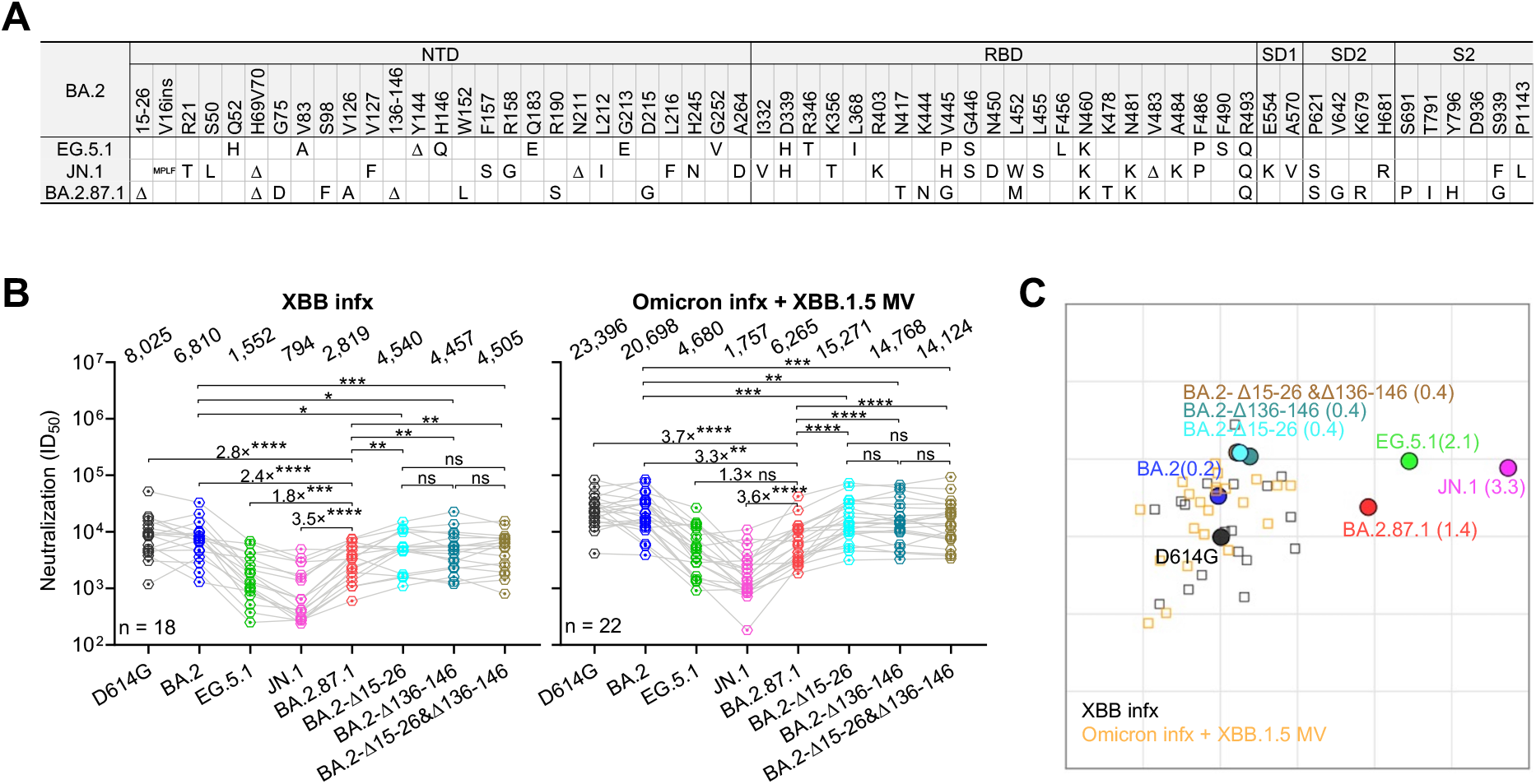
Spike mutations and serum neutralization of SARS-CoV-2 Omicron variant BA.2.87.1. **A**.Spike mutations of BA.2.87.1 in comparison of EG.5.1 and JN.1 on top of BA.2. Δ, deletion; ins, insertion. **B**.Neutralizing ID_50_ titers of serum samples from “XBB breakthrough” and “Omicron infx + XBB.1.5 MV” cohorts against the indicated SARS-CoV-2 variants. The geometric mean ID_50_ titers (GMT) are presented above symbols. Fold changes in neutralizing ID_50_ titers between BA.2.87.1 and other SARS-CoV-2 variants are denoted. Statistical analyses were performed by employing Wilcoxon matched-pairs signed-rank tests. ns, not significant; \**p*< 0.05; \*\**p* < 0.01; \*\*\**p* < 0.001; \*\*\*\**p* < 0.0001. n, sample size. Participants from the two cohorts all had received 3-4 doses of the wildtype monovalent vaccines, one dose of the BA.5 bivalent vaccine booster and then followed by either a XBB breakthrough infection (“XBB breakthrough”) or Omicron infection and XBB.1.5 monovalent vaccine booster (“Omicron infx + XBB.1.5 MV”). **C**.Antigenic map based on the neutralizing data from all serum samples in panel **B**. D614G represents the central reference for all serum cohorts, with the antigenic distances calculated by the average divergence from each variant. One antigenic unit (AU) represents an approximately 2-fold change in ID50 titer. Variant positions are shown as circles, while sera are denoted as gray and orange squares.

We accomplished this characterization by first assessing the serum virus-neutralizing titers in 43 participants from two distinct clinical cohorts: 1) “XBB infx,” comprising individuals who had an infection with the XBB sublineage virus; and 2) “Omicron infx + XBB.1.5 MV,” consisting of individuals who were administered an XBB.1.5 MV booster following a prior Omicron infection. All the participants had previously received 3-4 doses of the wildtype monovalent mRNA vaccines and 1 dose of the BA.5 bivalent vaccine booster. Detailed clinical information for each participant is provided in **Table S1**. Serum samples were collected on average 27.7 days post-infection or 25.5 days post-vaccination. We then determined their neutralizing antibody (NAb) titers using a pseudovirus neutralization assay against the ancestral D614G strain and several Omicron subvariants, BA.2, EG.5.1, JN.1, and BA.2.87.1, as well as BA.2 with added NTD deletions from BA.2.87.1: BA.2-Δ15-26, BA.2-Δ136-146, and BA.2-Δ15-26&Δ136-146 (**Figure 1B**). Our data suggested that both cohorts exhibited similar patterns of robust neutralization activity against all viruses tested, with the highest titers against D614G but substantially lower titers against the Omicron subvariants, particularly the currently dominant JN.1. This was despite the “Omicron infx + XBB.1.5 MV” cohort having approximately 3-fold higher neutralizing titers overall, compared to the XBB infection cohort. Notably, BA.2.87.1 showed roughly a 3.5-fold and a 1.5-fold higher neutralization susceptibility than JN.1 and EG.5.1, respectively, for both cohorts. Antigenic cartography based on these results revealed the BA.2.87.1 variant displayed an antigenic distance of 1.4 units from D614G, markedly less than that observed for JN.1, and less than that of EG.5.1, the prevalent variant in the preceding infection wave (**Figure 1C**). Next, an ACE2 inhibition assay and infectivity assay suggested that BA.2.87.1 did not exhibit more efficient engagement with the ACE2 receptor or higher entry efficiency into Vero-E6 and Calu-3 lung cells compared to EG.5.1 and JN.1 (**Figure S2 and S3**). Lastly, while the NTD deletions (Δ15-26, Δ136-146, or both) added to BA.2 conferred a subtle increase (∼1.5-fold) in serum antibody evasion, they did have a negative impact on virus infectivity (**Figure 1B, Figure S2 and S3**).

In summary, our findings suggest that the Omicron subvariant BA.2.87.1 is less immune-evasive and does not demonstrate more substantial engagement with viral receptor or infectivity compared to EG.5.1 and JN.1. Our observations are in line with results reported by Zhang et al. using sera from participants from Germany^2^; by Lasrado et al. using sera from participants in the US^3^; and by Yang et al., using sera from individuals in China^4^. Yet, they contrast with the findings of Wang et al. using sera from participants from China^5^. The causes of this inconsistency remain obscure, and the clinical relevance is uncertain. Nevertheless, our data may explain why BA.2.87.1 has not become more prevalent globally, given its virological characteristics, and they support the announcement by the USA CDC that the public health risk posed by BA.2.87.1 is expected to be low.

## Notes

### Competing Interest Statement

D.D.H. co-founded TaiMed Biologics and RenBio, acts as a consultant for WuXi Biologics and Brii Biosciences, and holds a director position on the board of Vicarious Surgical. A.G. was a member of the scientific advisory board for Janssen Pharmaceuticals. The other authors have no competing interests to declare.

