## supplemental file for "SARS-CoV-2 Omicron BA.2.87.1 Exhibits Higher Susceptibility to Serum Neutralization Than EG.5.1 and JN.1"

**Supplementary Appendix**

**Table S1. Demographics of clinical cohorts.**

Clinical information for each participant, including demographic, vaccine and infection history, and sera details. infx, infection; MV, monovalent vaccine; BV, bivalent vaccine.

| Sample ID | Age | Gender | Ethnicity | Infection time (year & month) | Vaccination and infection history | WT MV | BA.5 BV | Omicron infx | XBB.1.5 MV | Days post vaccination/infection |
|---|---|---|---|---|---|---|---|---|---|---|
| XBB infx | | | | | | | | | | |
| G3-1 | 33 | Female | Not Hispanic or Latino | 2023.02 | Pfizer-Pfizer-Pfizer-Pfizer-XBB | 3 | 1 | XBB | 0 | 30 |
| G3-3 | 38 | Female | Not Hispanic or Latino | 2023.08 | Pfizer-Pfizer-Pfizer-Pfizer-XBB | 3 | 1 | XBB | 0 | 27 |
| G3-4 | 59 | Female | Not Hispanic or Latino | 2023.08 | Pfizer-Pfizer-Pfizer-Pfizer-Pfizer-XBB | 4 | 1 | XBB | 0 | 26 |
| G3-5 | 44 | Male | Not Hispanic or Latino | 2023.05 | Pfizer-Pfizer-Pfizer-Pfizer-Moderna-XBB | 4 | 1 | XBB | 0 | 30 |
| G3-6 | 54 | Female | Not Hispanic or Latino | 2023.09 | Pfizer-Pfizer-Pfizer-Pfizer-Pfizer-XBB | 4 | 1 | XBB | 0 | 28 |
| G3-7 | 77 | Male | Not Hispanic or Latino | 2023.09 | Pfizer-Pfizer-Pfizer-Pfizer-Pfizer-XBB | 4 | 1 | XBB | 0 | 22 |
| G3-8 | 38 | Female | Not Hispanic or Latino | 2023.03 | Pfizer-Pfizer-Moderna-Pfizer-XBB | 3 | 1 | XBB | 0 | 28 |
| G3-9 | 59 | Female | Not Hispanic or Latino | 2023.05 | Pfizer-Pfizer-Pfizer-Pfizer-Pfizer-XBB | 4 | 1 | XBB | 0 | 30 |
| G3-10 | 41 | Female | Not Hispanic or Latino | 2023.02 | Pfizer-Pfizer-Pfizer-Pfizer-XBB | 3 | 1 | XBB | 0 | 29 |
| G3-11 | 38 | Female | Not Hispanic or Latino | 2023.08 | Pfizer-Pfizer-Pfizer-Pfizer-XBB | 3 | 1 | XBB | 0 | 30 |
| G3-12 | 42 | Male | Not Hispanic or Latino | 2023.04 | Pfizer-Pfizer-Pfizer-Moderna-XBB | 3 | 1 | XBB | 0 | 28 |
| G3-13 | 38 | Female | Not Hispanic or Latino | 2023.08 | Pfizer-Pfizer-Pfizer-Pfizer-XBB | 3 | 1 | XBB | 0 | 24 |
| G3-14 | 56 | Female | Not Hispanic or Latino | 2023.09 | Pfizer-Pfizer-Pfizer-Pfizer-Pfizer-XBB | 4 | 1 | XBB | 0 | 29 |
| G3-15 | 53 | Female | Not Hispanic or Latino | 2023.02 | Pfizer-Pfizer-Pfizer-Pfizer-XBB | 3 | 1 | XBB | 0 | 27 |
| G3-16 | 40 | Female | Not Hispanic or Latino | 2023.05 | Pfizer-Pfizer-Pfizer-Pfizer-XBB | 3 | 1 | XBB | 0 | 28 |
| G3-18 | 59 | Female | Not Hispanic or Latino | 2023.04 | Moderna-Moderna-Moderna-Moderna-Pfizer-XBB | 4 | 1 | XBB | 0 | 24 |
| G3-19 | 51 | Female | Not Hispanic or Latino | 2023.05 | Moderna-Moderna-Moderna-Moderna-Pfizer-XBB | 4 | 1 | XBB | 0 | 29 |
| G3-20 | 54 | Female | Not Hispanic or Latino | 2023.08 | Pfizer-Pfizer-Pfizer-Pfizer-XBB | 3 | 1 | XBB | 0 | 29 |
| **Mean** | **48.6** | | | | | | | | | **27.7** |
| Omicron infx + XBB.1.5 MV | | | | | | | | | | |
| G5-1 | 41 | Female | Not Hispanic or Latino | 2022.09 | Moderna-Moderna-Moderna-Pfizer-Pre-XBB Omicron-Moderna | 3 | 1 | Pre-XBB Omicron | 1 | 25 |
| G5-2 | 61 | Female | Not Hispanic or Latino | 2022.04 | Pfizer-Pfizer-Pfizer-Moderna-Pre-XBB Omicron-Moderna | 3 | 1 | Pre-XBB Omicron | 1 | 34 |
| G5-4 | 49 | Male | Not Hispanic or Latino | 2022.01 | Pfizer-Pfizer-Pfizer-Pfizer-Pre-XBB Omicron-Pfizer | 3 | 1 | Pre-XBB Omicron | 1 | 22 |
| G5-5 | 67 | Female | Not Hispanic or Latino | 2022.07 | Pfizer-Pfizer-Pfizer-Pfizer-Moderna-Pre-XBB Omicron-Moderna | 4 | 1 | Pre-XBB Omicron | 1 | 29 |
| G5-6 | 52 | Female | Not Hispanic or Latino | 2022.04 | Pfizer-Pfizer-Pfizer-Pfizer-Pfizer-Pre-XBB Omicron-Pfizer | 4 | 1 | Pre-XBB Omicron | 1 | 25 |
| G5-7 | 48 | Female | Not Hispanic or Latino | 2022.01 | Pfizer-Pfizer-Pfizer-Moderna-Pre-XBB Omicron-Pfizer | 3 | 1 | Pre-XBB Omicron | 1 | 22 |
| G5-8 | 43 | Male | Not Hispanic or Latino | 2022.01 | Pfizer-Pfizer-Pfizer-Pfizer-Pre-XBB Omicron-Pfizer | 3 | 1 | Pre-XBB Omicron | 1 | 20 |
| G5-9 | 53 | Female | Not Hispanic or Latino | 2022.10 | Pfizer-Pfizer-Pfizer-Pfizer-Pre-XBB Omicron-Pfizer | 3 | 1 | Pre-XBB Omicron | 1 | 25 |
| G5-10 | 40 | Female | Not Hispanic or Latino | 2022.04 | Pfizer-Pfizer-Pfizer-Moderna-Pre-XBB Omicron-Moderna | 3 | 1 | Pre-XBB Omicron | 1 | 21 |
| G5-11 | 37 | Female | Not Hispanic or Latino | 2022.05 | Pfizer-Pfizer-Pfizer-Moderna-Pre-XBB Omicron-Moderna | 3 | 1 | Pre-XBB Omicron | 1 | 20 |
| G5-12 | 43 | Female | Not Hispanic or Latino | 2022.01 | Pfizer-Pfizer-Pfizer-Moderna-Pre-XBB Omicron-Pfizer | 3 | 1 | Pre-XBB Omicron | 1 | 32 |
| G5-13 | 36 | Male | Not Hispanic or Latino | 2022.08 | Pfizer-Pfizer-Pfizer-Moderna-Pre-XBB Omicron-Moderna | 3 | 1 | Pre-XBB Omicron | 1 | 30 |
| G5-14 | 59 | Female | Not Hispanic or Latino | 2022.04 | Pfizer-Pfizer-Pfizer-Pfizer-Pfizer-Pre-XBB Omicron-Moderna | 4 | 1 | Pre-XBB Omicron | 1 | 29 |
| G5-15 | 35 | Female | Not Hispanic or Latino | 2022.06 | Pfizer-Pfizer-Pfizer-Pfizer-Pre-XBB Omicron-Pfizer | 3 | 1 | Pre-XBB Omicron | 1 | 29 |
| G6-1 | 54 | Female | Not Hispanic or Latino | 2023.02 | Pfizer-Pfizer-Pfizer-Moderna-Pfizer-XBB-Moderna | 4 | 1 | XBB | 1 | 22 |
| G6-2 | 58 | Female | Not Hispanic or Latino | 2023.02 | Pfizer-Pfizer-Pfizer-Pfizer-XBB-Pfizer | 3 | 1 | XBB | 1 | 22 |
| G6-3 | 61 | Male | Not Hispanic or Latino | 2023.05 | Pfizer-Pfizer-Moderna-Moderna-XBB-Moderna | 3 | 1 | XBB | 1 | 30 |
| G6-4 | 42 | Female | Not Hispanic or Latino | 2023.03 | Pfizer-Pfizer-Pfizer-Pfizer-Moderna-XBB-Pfizer | 4 | 1 | XBB | 1 | 29 |
| G6-5 | 62 | Female | Not Hispanic or Latino | 2023.06 | Pfizer-Pfizer-Pfizer-Pfizer-Pfizer-XBB-Pfizer | 4 | 1 | XBB | 1 | 21 |
| G6-6 | 46 | Male | Not Hispanic or Latino | 2023.05 | J&J-J&J-Moderna-Pfizer-XBB-Pfizer | 3 | 1 | XBB | 1 | 24 |
| G6-7 | 45 | Female | Not Hispanic or Latino | 2023.08 | Pfizer-Pfizer-Pfizer-Pfizer-XBB-Pfizer | 3 | 1 | XBB | 1 | 22 |
| G6-8 | 62 | Female | Not Hispanic or Latino | 2023.04 | Pfizer-Pfizer-Pfizer-Pfizer-Pfizer-XBB-Moderna | 4 | 1 | XBB | | 29 |
| **Mean** | **49.7** | | | | | | | | | **25.5** |

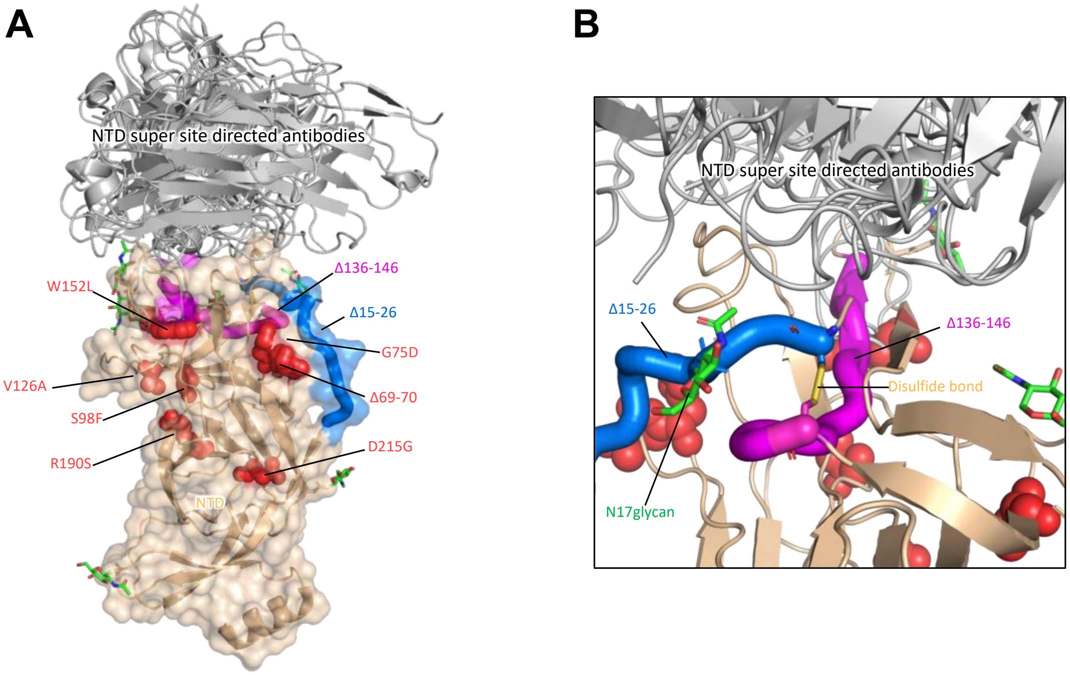

**Figure S1. Structural interpretation of NTD mutations in BA.2.87.1.**

A. Alterations in the NTD of the BA.2.87.1 variant. The deletion of amino acids 15-26 is shown using a marine-colored ribbon, while the deletion of amino acids 136-146 is illustrated with a magenta ribbon. Additional mutations within the NTD are indicated by red spheres.

B. The detailed interaction between the deletions at positions 15-26 and 136-146. The disulfide bond formation between cysteine residues at positions 15 and 136 is represented in orange sticks. The green sticks show the glycans on NTD. NTD, N-terminal domain;

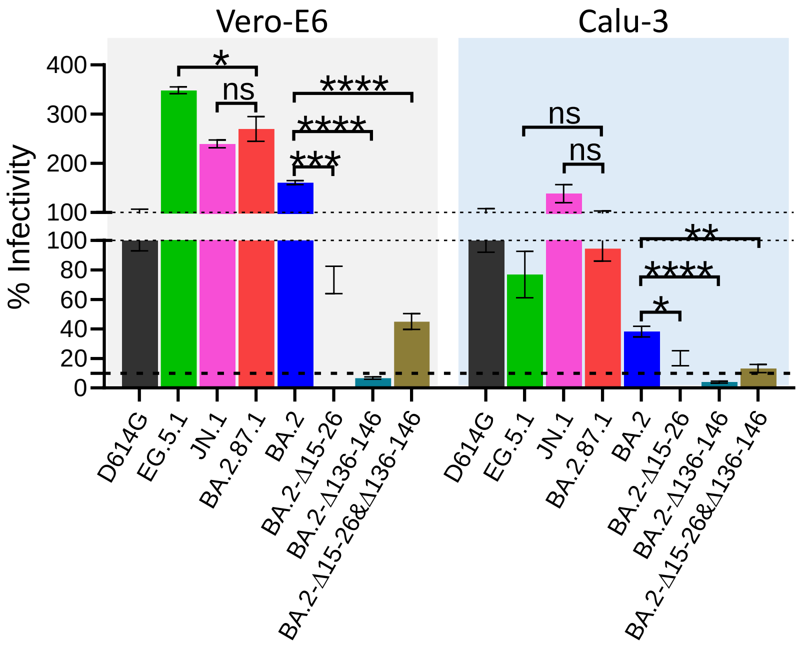

**Figure S2. Relative infectivity of pseudotyped SARS-CoV-2 viruses compared to D614G on Vero-E6 and Calu-3 cells.** Statistical analyses were performed using unpaired t tests. ns, not significant; *p < 0.05; **p < 0.01; ***p < 0.001; ****p < 0.0001

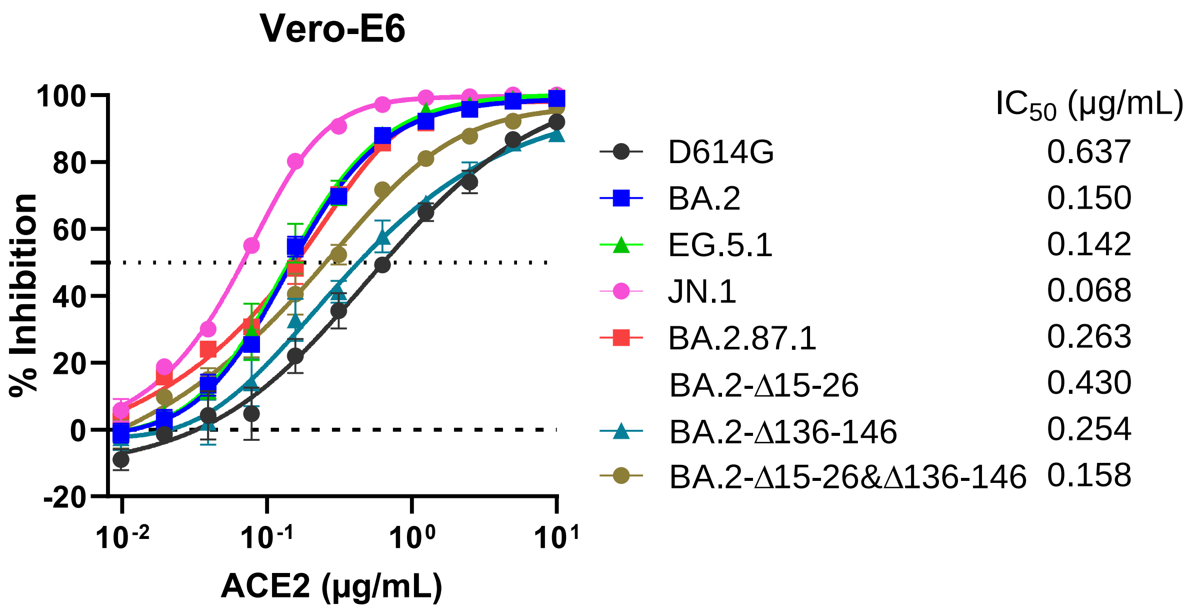

**Figure S3. Susceptibility of BA.2.87.1 pseudovirus to hACE2 inhibition compared to other SARS-CoV-2 variants on Vero-E6 cell with IC50 values denoted.**

**Supplementary Methods**

*Clinical cohorts*

Serum samples were collected as part of the Immunity Associated with SARS-CoV-2 Study (IASO), which was initiated in 2020 at the University of Michigan, Ann Arbor, Michigan[1]. Specimens were obtained following participant consent and in adherence to the protocol approved by the Institutional Review Board of the University of Michigan Medical School. Participants completed weekly symptom surveys, and if any symptoms were reported, they were tested for SARS-CoV-2. To detect potential breakthrough infections throughout the study, we analyzed all serum samples using the anti-nucleoprotein (NP) ELISA.

In this study, serum samples were collected from two cohorts: 1) individuals with a recent XBB sublineage infection who had not received the updated XBB.1.5 monovalent booster ("XBB infx"); 2) individuals who had recovered from an Omicron infection and had been administered the updated XBB.1.5 monovalent booster ("Omicron infx + XBB MV"). Participants from both cohorts were vaccinated with three or four doses of the original monovalent vaccine followed by one ancestral/BA.5 bivalent booster. The majority of the study subjects were female, representing 80%, with an average age of 49.2 years. Serum samples were collected, on average, 25.5 days post-XBB.1.5 vaccination and 27.7 days post-XBB sublineage infection. Demographic details, vaccination statuses, and serum collection timelines are detailed for each cohort in **Table S1**.

*Cell lines*

HEK293T (ATCC, CRL-3216) cells and Vero-E6 cells (ATCC, CRL-1586) were cultivated in Dulbecco's modified Eagle's medium (DMEM) supplemented with 10% heat-inactivated fetal calf serum and 1% penicillin-streptomycin. Calu-3 (ATCC,HTB-55) cells were maintained in Eagle's Minimum Essential Medium (MEM) supplemented with the same concentrations of heat-inactivated fetal calf serum and penicillin-streptomycin. All cell lines were cultured in an atmosphere of 5% $CO_2$ at 37°C.

*Construction of SARS-CoV-2 spike plasmids*

The spike gene of the BA.2.87.1 variant, as well as spike gene constructs bearing only large deletions (Δ15-26, Δ136-146, or both) in the N-terminal domain (NTD) found in BA.2.87.1, were generated by MEGAA[2] as previously described[3]. Briefly, 36 oligos used in MEGAA mutagenesis were synthesized from a SYNTAX STX-200 (DNA Script) with a high-fidelity run kit or sourced from Integrated DNA Technologies. The corresponding mutations were generated in the BA.2 spike gene construct using annealing, extension, ligation, and PCR steps. To confirm the sequences of the variants, we performed Illumina Miseq Nextera and Oxford Nanopore sequencing (MinION and R10.4.1), and analyzed sequencing data with Cutadapt 4[4], SAMtools 1.16.1 and BCFtools 1.17[5], fastp 0.23.4[6], bwa 0.7.17 (https://arxiv.org/abs/1303.3997), pod5 0.3.6 (Oxford Nanopore Technologies), dorado 0.5.3 (Oxford Nanopore Technologies), porechop 0.2.4[7], and minimap2 2.26[8]. All constructs were confirmed by visually inspecting sequencing data in IGV[9] and by Sanger sequencing.

*Pseudovirus production and infectivity*

Pseudotyped SARS-CoV-2 was produced following a previously established protocol[10]. HEK293T cells were first transfected with spike-encoding plasmids using 1 mg/mL of PEI and cultured for 24 hours. The transfected HEK293T cells were then infected with VSV-G pseudotyped ΔG-luciferase virus (Kerafast, EH1020-PM) at a multiplicity of infection (MOI) of approximately 3 to 5. Two hours later, the cells were washed three times with complete culture medium and cultured in fresh medium for another 24 hours. The transfection supernatant was then harvested and clarified by centrifugation at 2000 rpm for 10 minutes. Each viral stock was subsequently incubated with 20% I1 hybridoma (ATCC, CRL-2700) supernatant for 1 hour at room temperature to neutralize contaminating VSV-G particle before measuring titers and making aliquots for storage at −80°C until use.

In order to investigate virus entry host cell efficiency, pseudovirus particles bearing various SARS-CoV-2 spike proteins were inoculated onto Vero-E6 cells or Calu-3, starting with a volume of 50 µl per well in 96-well plates and followed by serial dilutions. After a 16-18 hour incubation at 37°C, the activity of virus-encoded firefly luciferase in cell lysates was measured. Subsequently, the luciferase activity for each tested pseudotyped virus, which was not oversaturated, was normalized to the parental D614G S control, with an infectivity set to 1.0.

*Pseudovirus neutralization assay and ACE2 inhibition assay*

Pseudoviruses were subjected to titration to standardize the viral input prior to each neutralization or inhibition assay. For neutralization assays, serum samples were inactivated at 56°C for 30 minutes before use, and the inactivated sera were diluted by a factor of 50 in a series of seven serial dilutions. For ACE2 inhibition assays, as reported in our study[10,11], soluble chimeric human ACE2, which contains ACE2 residues 1-732 and is fused to human IgG1 Fc, was diluted from 10 µg/mL with a dilution factor of two across 11 serial dilutions. Following this, pseudoviruses were added and incubated with varying dilutions of either the serum or ACE2 at 37 °C for 1 hour. As a control, wells containing only the pseudovirus were also prepared on each test plate. Subsequently, Vero-E6 cells were seeded at 40,000 cells per well and were incubated overnight at 37°C on either neutralization or inhibition test plates Afterwards, cellular lysis was conducted, and the resultant luciferase activity was quantified employing the Luciferase Assay System (Promega) in tandem with the SoftMax Pro v.7.0.2 software (Molecular Devices), in accordance with the manufacturer's instructions.The serum dilution that inhibits 50% of virus entry ($ID_{50}$), or the half-maximal inhibitory concentration ($IC_{50}$) by ACE2, was calculated using nonlinear five-parameter dose-response curve fitting using GraphPad Prism v.9.2.

*Antigenic cartography*

Antigenic cartography for the D614G variant and other SARS-CoV-2 variants was conducted through the integration of $ID_{50}$ titers from individual sera, as previously described[12]. The visual representations were generated utilizing the Racmacs package (version 1.1.4, accessible at https://acorg.github.io/Racmacs/) within the R computational environment, version 4.0.3. The algorithmic optimization process was executed over 2,000 iterations, with the 'minimum column basis' parameter explicitly set to 'none'. The computation of antigenic distances between each serum sample and corresponding variant was facilitated by the 'mapDistances' function. The D614G variant served as the center for the antigenic positioning of sera within each group, with the initial positions for the antigenic landscapes being manually adjusted to ensure accurate orientation of the JN.1 variant in relation to D614G. To maintain uniformity across the antigenic landscapes, all maps were aligned with the D614G variant situated at the central left position, thereby establishing a consistent point of reference throughout the study.

*Quantification and statistical analysis*

$ID_{50}$ values for serum neutralization and $IC_{50}$ values for ACE2 inhibition were obtained from a five-parameter dose-response curve using GraphPad Prism v.9.2. Statistical analyses were conducted by either Wilcoxon matched-pairs signed-rank tests or unpaired t tests in the same software, as indicated in figure legends. Levels of statistical significance are annotated as: ns, not significant; *, $p < 0.05$; **, $p < 0.01$; ***, $p < 0.001$ and ****, $p < 0.0001$.

### Acknowledgements

This research received financial support from the NIH SARS-CoV-2 Assessment of Viral Evolution (SAVE) Program (Subcontract No. 0258-A709-4609 under Federal Contract No. 75N93021C00014) and the Gates Foundation (project INV019355) to D.D.H., as well as funding from the NIH contract 75N93019C00051 to A.G.  We also acknowledge funding support from the NSF (MCB-2032259) and R21 (1R21HG013161) attributed to H.H.W. We thank all who have submitted data to GISAID. We would like to extend our thanks to Liyuan Liu, and Yiming Huang for their assistance with plasmid construct generation and experimental insight. Furthermore, we express our gratitude to Zijin Chu, Theresa Kowalski-Dobson, Carmen Gherasim, David Manthei, Anna Buswinka, Gabe Simjanovski, Joseph Wendzinski, Mayurika Patel, Kathleen Lindsey, Dawson Davis, Victoria Blanc, Savanna Sneeringer, and Pamela Bennett-Baker of the IASO study team for conducting the IASO study.

### Author Contributions

L. L. and D.D.H. conceived and supervised this project.  Q.W. managed the project.  Q.W., I.A.M., and L. L. conducted experiments. L.T.S., Y.S., Y.Q., and H.H.W. constructed the spike expression plasmids. Y.G. conducted bioinformatic analyses. E.S., R.V., and A.G. provided clinical samples. Q.W., I.A.M., Y.G., L. L., and D.D.H. analyzed the results. Q.W., I.A.M., Y.G., L.T.S., H.H.W., L. L., ,D.D.H. wrote the manuscript. All authors reviewed the results and approved the final version of the manuscript.

### Declaration of Interests

D.D.H. co-founded TaiMed Biologics and RenBio, acts as a consultant for WuXi Biologics and Brii Biosciences, and holds a director position on the board of Vicarious Surgical. A.G. was a member of the scientific advisory board for Janssen Pharmaceuticals. The other authors have no competing interests to declare.
